# Personal Cancer Genome Reporter: variant interpretation report for precision oncology

**DOI:** 10.1101/122366

**Authors:** Sigve Nakken, Ghislain Fournous, Daniel Vodák, Lars Birger Aasheim, Ola Myklebost, Eivind Hovig

## Abstract

**Summary:** Individual tumor genomes pose a major challenge for clinical interpretation due to their unique sets of acquired mutations. There is a general scarcity of tools that can *i)* systematically interrogate cancer genomes in the context of diagnostic, prognostic, and therapeutic biomarkers, *ii)* prioritize and highlight the most important findings, and *iii)* present the results in a format accessible to clinical experts. We have developed a stand-alone, open-source software package for somatic variant annotation that integrates a comprehensive set of knowledge resources related to tumor biology and therapeutic biomarkers, both at the gene and variant level. Our application generates a tiered report that will aid the interpretation of individual cancer genomes in a clinical setting.

**Availability and Implementation:** The software is implemented in Python/R, and is freely available through Docker technology. Documentation, example reports, and installation instructions are accessible via the project GitHub page: https://github.com/sigven/pcgr)

## 1 Introduction

High-throughput molecular profiling of tumor genomes can improve clinical decision support of individual cancer cases. Although detection of acquired genetic alterations in tumors is still a challenging matter, we and others would argue that the major bottleneck for precision oncology occurs downstream of variant calling (Good *et al.*, 2014). Tumor genome analyses will only be useful for oncologists when the identified somatic variants can be prioritized based on comprehensive integration with existing knowledge on clinical markers that highlights potential therapeutic options.

A range of different tools have been developed for functional annotation of genomic variants (Wang *et al.*, 2010; Ramos *et al.*, 2015; McLaren *et al.*, 2016). Although some have a specific focus on the oncology domain (e.g. *Oncotator*), they are all largely targeted towards the research community. Existing solutions offer limited support for summaries and reports at the level of individual cancer genomes, particularly when it comes to clinical relevance. Moreover, the degree of quality control and update frequency of annotation resources vary considerably. Recently, we have seen significant development of databases that harvest reports from the scientific literature about cancer genome variants and their particular relationships to tumorigenesis, druggability, and clinical outcomes. These include the Database of Curated Mutations (DoCM), The Drug Gene Interaction Database (DGIdb), and most importantly the community-driven database of Clinical Interpretations of Variants in Cancer (CIViC)(Ainscough *et al.*, 2016; Wagner *et al.*, 2016; Griffith *et al.*, 2017). Other resources covering known cancer mutation hotspots, mutational signatures, and predicted driver mutations have also emerged, which collectively indicate potential underlying mechanisms of tumor development and relevance for different treatment regimes (Petljak and Alexandrov, 2016; Chang *et al.*, 2016; Secrier *et al.*, 2016; Gonzalez-Perez *et al.*, 2013).

As part of a national collaboration on personalized cancer medicine (Norwegian Cancer Genomics Consortium), we have developed the Personal Cancer Genome Reporter (PCGR), a software package for the generation of clinically interpretable reports of individual cancer genomes. The software extends basic variant annotations from Variant Effect Predictor (VEP) with oncology-relevant annotations retrieved flexibly with *vcfanno* (Pedersen *et al.*, 2016), and produces HTML reports that can be navigated by clinical oncologists.

## 2 PCGR workflow

The pipeline for generation of personal cancer genome reports comprises four major steps: (1) basic variant consequence annotation using VEP, (2) allele-specific annotation for precision oncology using *vcfanno*, (3) functional and cancer-focused gene annotation, and (4) estimation of mutational signature contributions, summary, prioritization and reporting with the R language and R markdown templates (Figure 1). All software components are integrated and provided by means of the Docker technology, implying that all underlying dependencies are packaged into a standardized software container. The Docker solution was chosen to offer individual labs a simple installation of PCGR in their in-house workflow for high-throughput analysis of tumor genomes. The application comes with an annotation data bundle, which we plan to update on a quarterly basis. The human genome assembly GRCh37 is currently supported.

**Figure 1:**
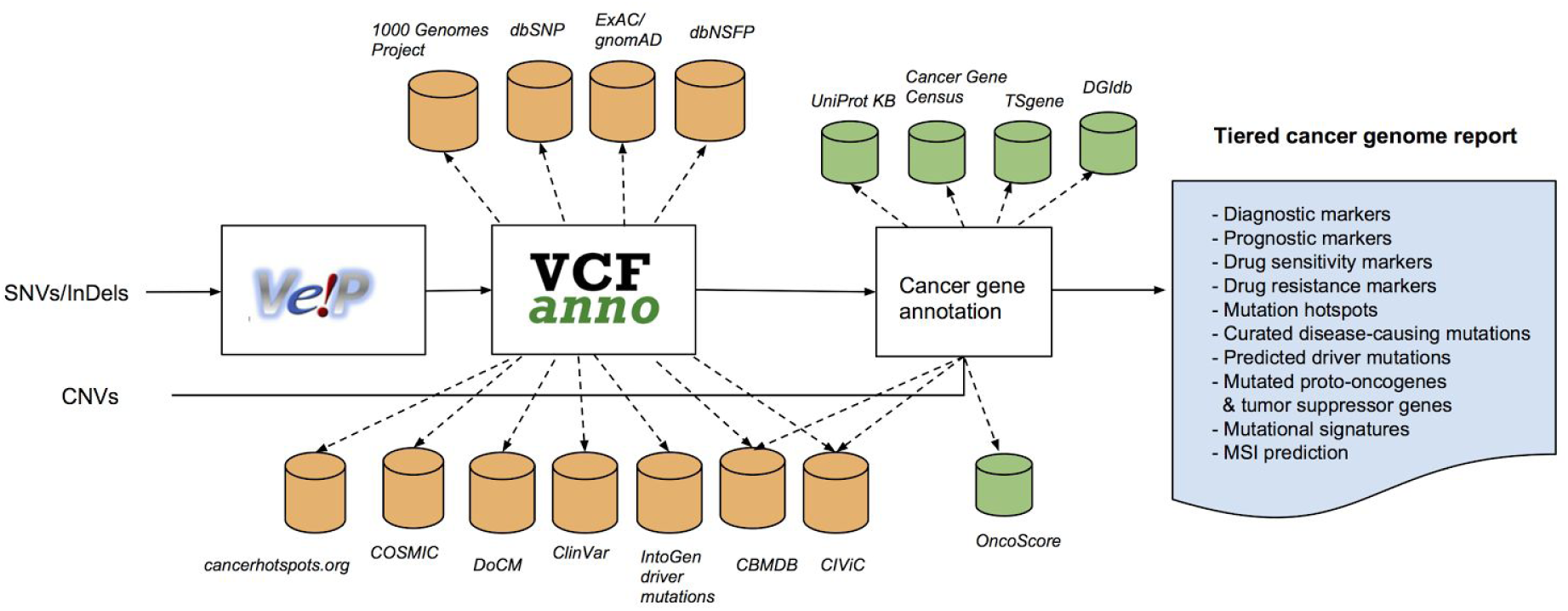
PCGR workflow. Abbreviations: CNV = Copy number aberration, SNV = Single nucleotide variant, InDel = Insertion/deletion, dbNSFP = database of non-synonymous functional predictions, COSMIC = Catalogue of Somatic Mutations in Cancer, DoCM = Database of Curated Mutations, CIViC = Clinical Interpretations of Variants in Cancer, ExAC = Exome Aggregation Consortium, gnomAD = Genome Aggregation Database, DGIdb = Drug Gene Interaction Database, dbSNP = Database of Short Genetic Variations, TSgene = Tumor suppressor gene database, CBMDB = Cancer bioMarkers Database

The PCGR workflow accepts two types of input files: 1) a single-sample VCF file encoding the genomic coordinates of somatic SNVs/indels, and 2) a basic somatic copy number segment file that encodes chromosomal segment locations and their log-2 ratios (specific requirements are given in the GitHub documentation).

For SNVs and indels encoded in a VCF file, VEP is utilized to determine variant consequence information. For the sake of simplicity and ease of use, the VEP annotation is run with a fixed set of parameters that includes all gene cross-references, protein domain annotations, and overlap with regulatory regions, using GENCODE as the underlying gene transcript model. Check for co-located known variants is omitted in VEP and instead undertaken by *vcfanno* to ensure up-to-date and more comprehensive cancer-relevant annotations. All transcript-dependent consequences per variant are retained in the VEP-annotated VCF file, and the consequence block of highest functional relevance (as provided by VEP’s *--flag_pick* option) is flagged for further downstream analysis. Next, *vcfanno* is applied on multiple variant databases in parallel in order to enrich each somatic call with allele-specific annotations that are directly or indirectly relevant for clinical interpretation (as depicted in Figure 1). These allele-specific annotations include *i)* population-specific germline allele frequencies (for quality control and potential filtering), *ii)* pathogenicity predictions for splice-site and missense variants by multiple algorithms, *iii)* overlap with known mutational cancer hotspots and previously predicted driver mutations, *iv)* tissue/tumor type frequency in case of previously detected somatic variants, *v)* known disease/cancer associations, and *vi)* clinical evidence items of relevance for diagnosis, prognosis, or drug sensitivity/resistance.

In the third step of the workflow, gene-level annotations are aggregated. Known proto-oncogenes and tumor suppressors are marked, as are other genes causally implicated (either by prediction or curation) in the development of different tumor types. Antineoplastic agents and their molecular targets are also annotated. Each gene is finally assigned a score that reflects its relative strength of association to cancer in the biomedical literature, enabling ranking of novel variants according to functional relevance (Rocco *et al.*, 2017). The annotation of copy number aberrations is limited to this third step, in which gene transcripts that intersect gained or depleted segments are identified, and clinical and etiologic cancer associations are retrieved.

The final and fourth step of the PCGR workflow summarises and prioritizes the annotated variants in a structured and interactive report, adopting recently proposed recommendations (Ritter *et al.*, 2016; Dienstmann *et al.*, 2014). Specifically, a tiered report is constructed, starting from actionable markers in *Tier 1,* toward aberrations relevant for tumorigenesis in *Tier 2 and 3*, and ending with variants of unknown functional relevance in *Tier 4 and 5*. In addition to the tier structure, mutated genes in *Tier 3-5* are prioritized by means of a literature-derived score for oncogenic potential, which draws attention to the most relevant findings. Finally, the report offers optional prediction of microsatellite instability, in addition to estimates of known mutational signatures present in the tumor, and their associated underlying etiologies (Rosenthal *et al.*, 2016).

## 3 Discussion

We have utilized the Docker technology to develop a stand-alone report engine for clinical interpretation of cancer genomes. The tool has a particular focus on coding variants, i.e. variants found through exome sequencing. A limitation of the current implementation is that actionable aberrations are not taking into account the tissue/tumor type of the query, all markers of clinical utility are now being reported. Moreover, we foresee that the report can be significantly strengthened through the addition of other molecular profiling datasets, such as gene expression. Expression will not only add an important layer on top of results found at the DNA level, it can also add significant value towards assessment of therapeutic potential by other analyses, such as inference of cell type composition in the tumor tissue, and *insilico* predictions of drug response (Newman *et al.*, 2015; Geeleher *et al.*, 2014).

